# Medical Grade Manuka Honey Inhibits A23187 Induced Mast Cell Degranulation and Cytokine Release via Downregulation of the ERK Signalling Pathways

**DOI:** 10.64898/2026.08.02.742303

**Authors:** K Garba, OKA Abdelwahab, LC Lau, ME Khedr, M Yusuh, AF Walls, B Birch, BA Lwaleed

**Affiliations:** School of Health Sciences, Faculty of Environmental and Life Sciences, University of Southampton, UK; Faculty of Medicine, University of Southampton, UK

**Keywords:** Manuka honey, mast cell, degranulation, cytokines, MAPK, signalling

## Abstract

**Rationale:** Mast degranulation is a driver of several pathologies. Several endogenous peptides have been established as stimulators of mast cell degranulation. Compounds that stabilise mast cells have in part demonstrated inhibitory effect on the inflammatory cascade. Manuka honey has been widely used for the treatments of various inflammatory diseases. However, its effect on A23187 (Calcium ionophore) stimulated mast cell degranulation, cytokine release and cellular signalling pathways has not yet been investigated.

**Aim of the study:** We aim to investigate the effect of Medical Grade Manuka Honey (MGMH) on A23187 induced mast cell degranulation, cytokine release and downstream signalling pathways in a human mast cell lines LAD2 model.

**Materials and methods:** The cytotoxic effect of MGMH on LAD2 cells was assessed using LDH assay. LAD2 cells were pre-incubated with MGMH and challenged with A23187.Mast cell degranulation was measured by β-hexosaminidase release whilst histamine and cytokines released were measured by ELISA. The effect of MGMH on downstream signalling of Mitogen Activated Protein Kinase (MAPK) pathways was determined and quantified by SDS-PAGE western blotting.

**Results:** MGMH at 2% and 4% was well tolerated by LAD2 cells. MGMH at all concentrations tested significantly inhibited the A23187 triggered release of β-hexosaminidase but failed to inhibit the release of histamine. MGMH 4% significantly inhibited the release of GM-CSF and IL-8. MGMH (2% and 4%) significantly down regulated the expression of both ERK I and ERK II. However, MGMH at all the doses tested had no effect on the expressions of JNK and p38. On the contrary, an increase expression of these p38 and JNK were noted with MGMH pre-incubation.

**Conclusion:** Our present study provides evidence that MGMH inhibition of A23187 stimulated degranulation of mast cells and cytokine release through down regulation of ERK I and II signalling. The results suggest potential use as a mast cell stabiliser and effective treatments of mast cell mediated inflammatory diseases.

**Impact:** This study provides the first evidence that Medical Grade Manuka Honey (MGMH) can attenuate A23187-induced mast cell activation and pro-inflammatory cytokine release in human LAD2 mast cells through modulation of ERK1/2 signalling pathways. The findings advance our understanding of the cellular and molecular mechanisms underlying the anti-inflammatory properties of Manuka honey.

The demonstration that MGMH inhibits β-hexosaminidase release suggests a mast cell-stabilising effect, highlighting its potential as a novel natural therapeutic agent for mast cell-mediated disorders, including allergic diseases, interstitial cystitis, asthma, chronic inflammatory skin conditions, and mast cell activation syndromes. The observed reduction in GM-CSF and IL-8 production further indicates that MGMH may help limit the amplification and persistence of inflammatory responses.

Mechanistically, the selective downregulation of ERK1/2 signalling provides new insight into how MGMH exerts its biological effects and identifies a potential molecular target through which its anti-inflammatory activity is mediated. These findings contribute to the growing evidence base supporting the therapeutic value of Manuka honey beyond its established antimicrobial and wound-healing properties.

Overall, this work lays the foundation for future preclinical and clinical studies investigating MGMH as a safe, naturally derived mast cell stabiliser and anti-inflammatory intervention, with potential applications across a broad spectrum of allergic and inflammatory diseases.

## INTRODUCTION

The degranulation of mast cells plays an important role in several inflammatory diseases such as interstitial cystitis (IC), neuropathic pain, migraine, cancer, atopic dermatitis, psoriasis, autoimmune disease and allergic conditions amongst others (Anupam et al. 2015; Lyons and Pullen 2020). The activation of mast cells to release mediators is regulated through IgE-dependent and IgE-independent mechanisms; the former drives Type-1 hypersensitivity reactions and allergic reactions whilst the latter underlies the basis for chronic inflammatory diseases (Tkaczyk and Gilfillan 2001; Tatemoto et al. 2006).

In humans, mast cells are anatomically positioned in vascularized tissues to provide first line defence against invading antigens (adaptive immunity) by releasing inflammatory mediators such as proteases, lysosomal enzymes, proteoglycans, biogenic amines and cytokines (Galli et al. 2005; Gri et al. 2012). Despite the defensive functions that mast cells provide, dysregulation of mast cell homeostasis, as seen in mastocytosis or mast cell activation syndrome, sets the stage for chronic inflammation (Akin 2017). This is due to the secretion of cytoplasmic and nuclear mediators that orchestrate the inflammatory process. Consequently, inhibiting mast cell degranulation could provide a pathway for the therapy of mast cell aberrant function.

Calcium ionophore (A23187) promotes the influx of calcium ions into the endoplasmic reticulum and via this mechanism acts as a non-immunologic stimulant of mast cell degranulation (Foreman et al. 1973; Wightman et al. 2002). This agent was used to stimulate the mast cell secretory functions in human LAD2 cells. The LAD2 cell line was chosen due to its SCF-dependency, consistent degranulation and ability to withstand long term storage without losing its critical phenotypic functions as opposed to the alternative cell line HMC-1 (Kirshenbaum et al. 2003).

Natural products and their derivatives provide an alternative route for drug discovery and disease treatment. Thus, they are seen to have an established role in the historical treatment of a plethora of disease conditions (Newman and Cragg 2016). As such, Medical Grade Manuka Honey (MGMH) is popular as a nutraceutical (as opposed to pharmaceutical) agent. It has demonstrated considerable anti-inflammatory, anti-microbial, anti-proliferative and anti-oxidant actions (Alvarez-Suarez et al. 2014; Burns et al. 2018). Manuka honey is obtained from the indigenous New Zealand plant, *Leptospermum scoparium* or Manuka Myrtle. It is rich in flavonols, phenols and volatile oils. Given the diverse involvement of mast cell degranulation in several inflammatory diseases and the reported anti-inflammatory actions of Manuka honey; we hypothesized that Manuka Honey could be a valuable lead compound in inhibiting mast cell degranulation. Thus, the aim of this work was to investigative the effects of Medical Grade Manuka Honey (MGMH) on A23187 induced mast cell degranulation, cytokine release and downstream signalling pathways in an in vitro model of human mast cell lines LAD2.

## MATERIALS AND METHODS

### Reagents

MGMH (Comvita New Zealand Ltd, New Zealand), Histamine ELISA kit (Neogene, UK), Triton X (Sigma), A13187 (Sigma), Cytotoxicity detection kit-LDH (Roche), RIPA buffer (Cell signalling Technology), Phenyl methyl sulphonyl fluoride (Sigma), beta actin Rabbit mAB (Cell signalling), Human p-ERK1/ERK2 rabbit mAB (R and D system), p38 rabbit mAB (Cell signalling Technology), anti-rabbit IgG HRP-linked (Cell signalling Technology), β-actin (Cell signalling Technology), 4-12% precast gels (Sigma), Bovine Serum Albumin (Sigma), running buffer (Sigma) and transfer buffer (Sigma).

### Cell culture

Laboratory of Allergic Disease 2 (LAD2) cells were kindly gifted by Professor Arnold Krishenbaum of the National Institute of Allergic Disease, USA. The cells were maintained in Stem pro-34, with growth medium (10 ml/l), supplemented with Stem Cell Factor (100 ng/ml), penicillin (100U/ml) and streptomycin (100U/ml). Cells were maintained at 70-80% confluency at a density of 0.5X10^6^ cells/ml by passaging after every 10 days using hemi-depletion.

### Cell viability assays

Approximately 5X10^4^ LAD2 cells were pre-treated with vehicle, 2%, 4%, 6% and 8% concentrations of MGMH for 30 minutes on 96-well cell culture plates in a humidified cell culture incubator at 37^0^C and 5% CO2. The vehicle served as negative control. After incubation, the cell suspension was washed with an equal volume of PBSX1 and the resultant cell suspension centrifuged at 5000 g at 4^0^ C for 5 minutes. The resultant cell pellets were resuspended in Tyrodes buffer. The background control consisted of Tyrodes buffer only in the wells; the low control was Tyrodes buffer and cell suspension in 1:1 ratio while the high control (+ control) was a cell suspension and 1% triton (final concentration) in 1:1 ratio. To the resuspended cells in Tyrodes buffer 100 µl of freshly prepared reaction mixture (LDH assay kit, Roche) was added and the plate incubated for 30 minutes while protected from light. The reaction is a colorimetric assay with a maroon colour developing in proportion to the amount of lactate dehydrogenase released. Absorbance was read at 490 nm and the reference wavelength was set at 595 nm. Net release was calculated by the formula below after correcting for background control in each case.

LDH net release= (sample-low control)/ (High control) x 100

### Degranulation assay

For calcium ionophore induced degranulation about 1X10^5^ LAD2 cells were pre-treated with Tyrodes buffer, MGMH (2%, 4%, final well concentration) for 30 minutes and challenged with A23187 for 40 minutes in 96 well V-bottom cell culture plates. Total β-hexosaminidase was obtained by lysing equal amounts of untreated cells with Triton X 1% in PBS (total). The plates were centrifuged at 225g for ten minutes at room temperature. 30 µl of the resulting supernatant was transferred to a 96-well assay plate. 50 µl of a β-hexosaminidase substrate (p-Nitrophenyl-N-Acetyl-β-D-glucosaminide, in 0.1M citrate buffer pH 4.5) was added to the supernatant for 60 minutes. The reaction was stopped by addition of 100 µl 0.1 M glycine. Absorbance was read at 410 nm on a plate reader.

The net release of β-hexosaminidase was calculated from the formula below:

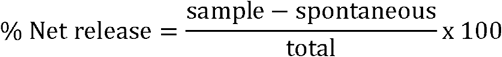

### Histamine ELISA assay

About 1X10^4^ LAD2 cells were pre-treated with Tyrodes buffer, MH (2% and 4%) for 30 minutes and challenged with A23187 0.3 µM for 40 minutes in 96 well V-bottom cell culture plates. The plates were centrifuged at 225 g for ten minutes at room temperature. About 50 µl of supernatant was used for the histamine ELISA assay according to the manufacturer’s instructions (Neogene, UK). The amount of Histamine released was read from the standard curve in ppm.

### Pro-inflammatory cytokines (IL-1β, GM-CSF, IL-8) ELISA assay

About 4X10^6^ LAD2 cells were pre-treated with Tyrodes buffer, MGMH (2%, 4%) for 30 minutes and challenged with A23187 0.3 µM for 24 hours in 96 well V-bottom cell culture plates. The plates were centrifuged at 225 g for ten minutes at room temperature. 25 µl of supernatant was used for the cytokine assay as described according to the manufacturer’s protocol (MSD Uplex Group 1 (Human), Meso Scale Discovery, USA).

### SDS-PAGE western blotting

About 1X10^6^ LAD2 cells were pre-treated with Tyrodes buffer, MGMH (2%, 4% and 6%) for 30 minutes and challenged with A23187 0.3 µM for 15 minutes. The reaction was stopped by adding equal amounts of cold PBSX1 and the cell suspension centrifuged for 5 minutes at 5000 g. Supernatants were discarded and cells were lysed with RIPA buffer in 1 M PMSF. Protein content was determined using a Pierce BCA protein assay (ThermoFisher). Equal amounts of protein (25 µg) were electrophoresed on 8-12% SDS precast gels and transferred to a nitrocellulose membrane. The membrane was blocked with 5% non-fat dry milk in Tris buffered saline Tween (TBST) for one hour at room temperature and washed three times each for five minutes. Blots were probed using primary antibodies with gentle shaking at 4°C overnight. The primary antibodies were: p-p38 (1:1000), p-JNK (1:1000) and p-ERK (0.5 µg/ml) and β-actin (1:1000). The membrane was washed three times with TBST for five minutes each and incubated with anti-rabbit IgG HRP-linked secondary antibody (1:2000) for one hour at room temperature. Blots were washed three times with TBST each for five minutes. Phosphorylated bound proteins were developed by use of an enhanced chemiluminiscence substrate (ThermoFisher, UK) for one minute, wrapped in photographic film and imaged using Chemidoc (Biorad, UK).

### Statistical analysis

Statistical analysis was performed using Graphpad prism 9.2.0 for Windows, San Diego, California, USA (www.graphpad.com). Data were analysed using one way ANOVA in comparison with A23187 group. This was followed by Dunnete post hoc test; p≤0.05 was considered significant.

## RESULTS

### The effect of MGMH on LAD2 cells

Pre-treatment of LAD2 cells with 2% and 4% MGMH for 30 minutes did not result in any cytotoxicity as shown in Figure 1. However, MGMH 6% and 8% at that exposure time led to a significant release of the LDH enzyme; suggesting cytotoxicity of these concentrations. Thus, MGMH 2% and 4% were used for degranulation, cytokine and SDS PAGE western blotting studies, as shown in figure 1.

**Figure 1:**
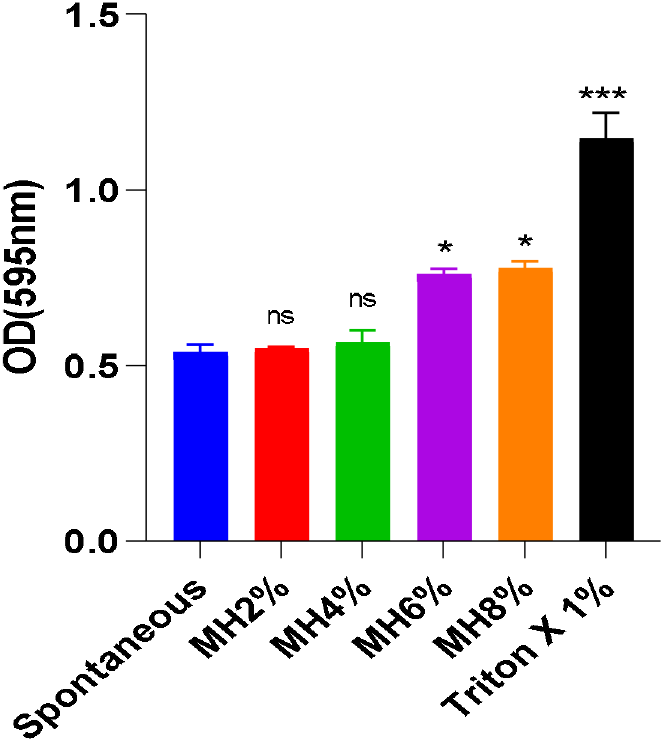
The effect of various concentrations of MGMH on the release of lactate dehydrogenase assay. LAD2 cells were seeded in 96-well plates, subsequently Tyrodes buffer (Spontaneous), 2%-8% MGMH, and 1%Triton X were added. The release of LDH was quantified. Data is expressed as mean OD ± SEM. A one way ANOVA was used to compare differences in concentrations and control followed by Dunette post hoc test. Value of P was set at ≤0.05. Experiment was done in triplicate and n=3. *≤0.05, ***=0.000, ns= non-significant.

### MGMH inhibited A23187 induced release of beta hexosaminidase but not histamine

The effect of MGMH on the release of β-hexosaminidase was investigated. At 2% and 4%, MGMH significantly inhibited the degranulation of mast cells. However, the same concentration failed to inhibit the release of histamine (Figure 2).

**Figure 2:**
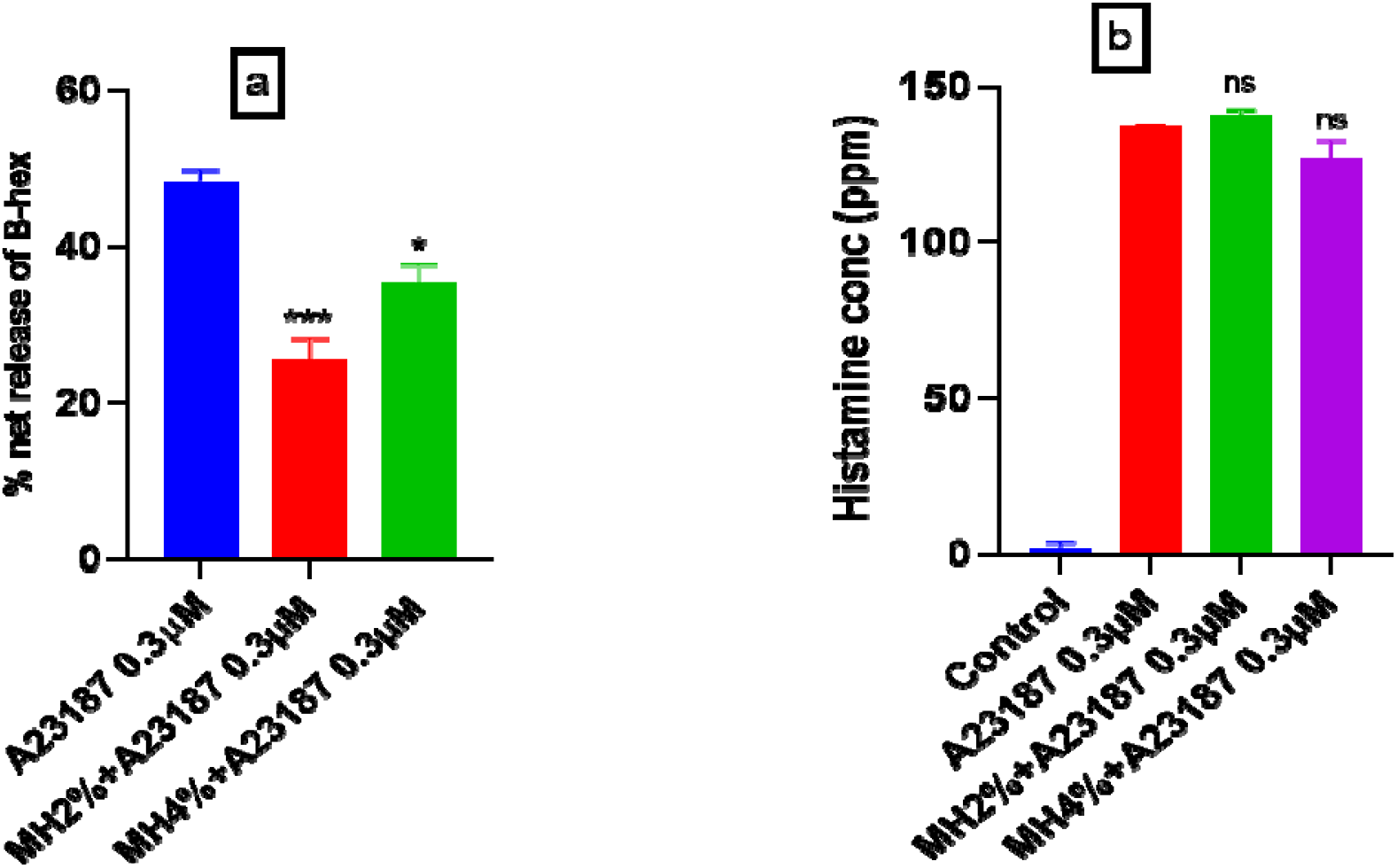
The effect of MGMH A23187 induce release of beta hexosaminidase and histamine. LAD2 cells were pre-treated with MGMH 2% and 4% for 30 minutes and the stimulated with A23187 0.3 µm for 40 minutes, β hexosaminidase release (a) and histamine release (b). Data is presented as Mean ± SEM of the % inhibition. A one way ANOVA was used to compare differences of treatments with SP group followed by Dunette post hoc test. Experiment was repeated three times. *≤0.05, ***=0.000, ns= non-significant.

### The effect of MGMH on cytokines release by LAD2 cells

To investigate the effect of MH on the A23187 mediated release of pro-inflammatory cytokines, ten cytokines were tested: TNF-α, MCP-1, IL-1β, IL-8, MIP1, IP10, ITAC, IL-17A, GRO and GM-CSF. However, only three cytokines were detected in the cell culture supernatant (Figure 3). The remaining cytokines concentrations were too low in the supernatant to be detected. 4% MGMH significantly inhibited the spontaneous and A23187 stimulated release of IL-8 and GM-CSF. Interestingly, MGMH 4% alone evoked the release of IL-1β without A23187 stimulation (Figure 3).

**Figure 3:**
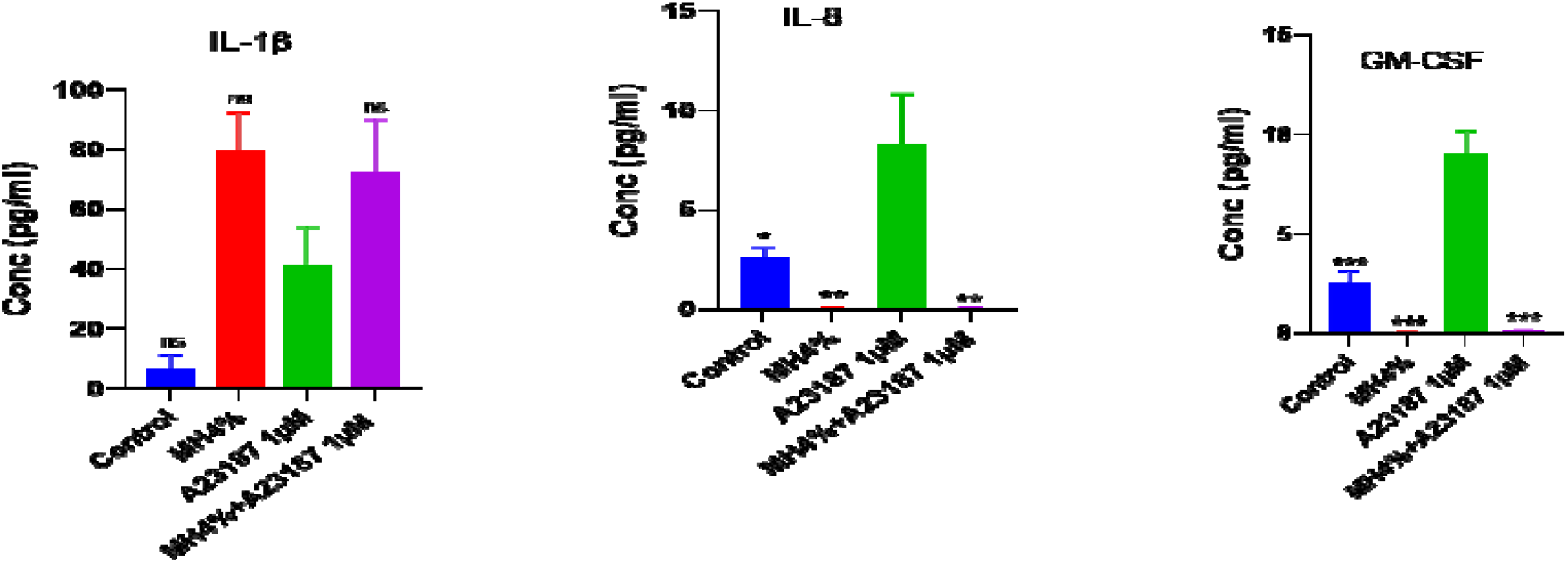
Effect of MGMH on A23187 induced release of cytokines. LAD2 cells were seeded in a 6-well plates with myeloma IgE for 24 hours. After which LAD2 cells were washed with PBSX1 and pre-incubated with 2% and 4% MGMH for 30 minutes, then stimulated with αIgE for 40 minutes. The release of β-hexosaminidase was quantified. Data is presented as Mean ± SEM of the % inhibition n=3. A one way ANOVA was used to compare differences of treatments with αIgE group followed by Dunette post hoc test. Experiment was repeated three times. **=0.001, ns= non-significant.

### The effect of MGMH on ERK expression by the LAD2 cells

Western blotting was performed to detect the effect of MGMH on p38, JNK and ERK (Figure 4). As shown in Figure 5, the results indicate that MGMH increased the expression of p38 and JNK. MGMH at concentrations at 2% and 4% inhibited the expression of ERK I in a concentration dependent manner. A similar effect was observed with respect to the expression of ERK II.

**Figure 4:**
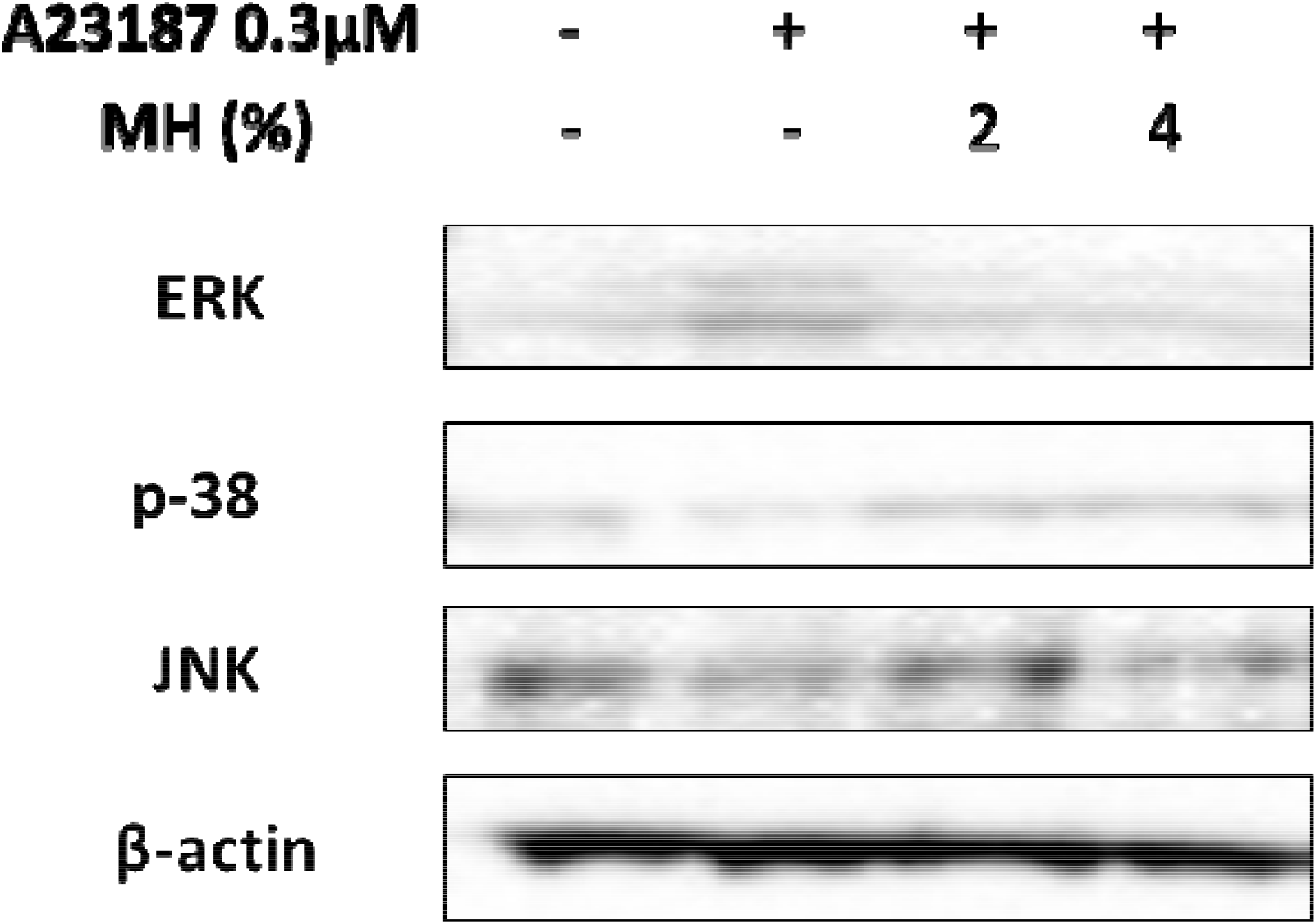
An immunoblot showing the effect of MH on A23187 induced expressions of signalling proteins. LAD2 cells were pre-incubated with 2% and 4% MGMH and stimulated with A23187 0.3 µm. Activation of ERK, P38 and JNK were assessed using western blotting.

**Figure 5:**
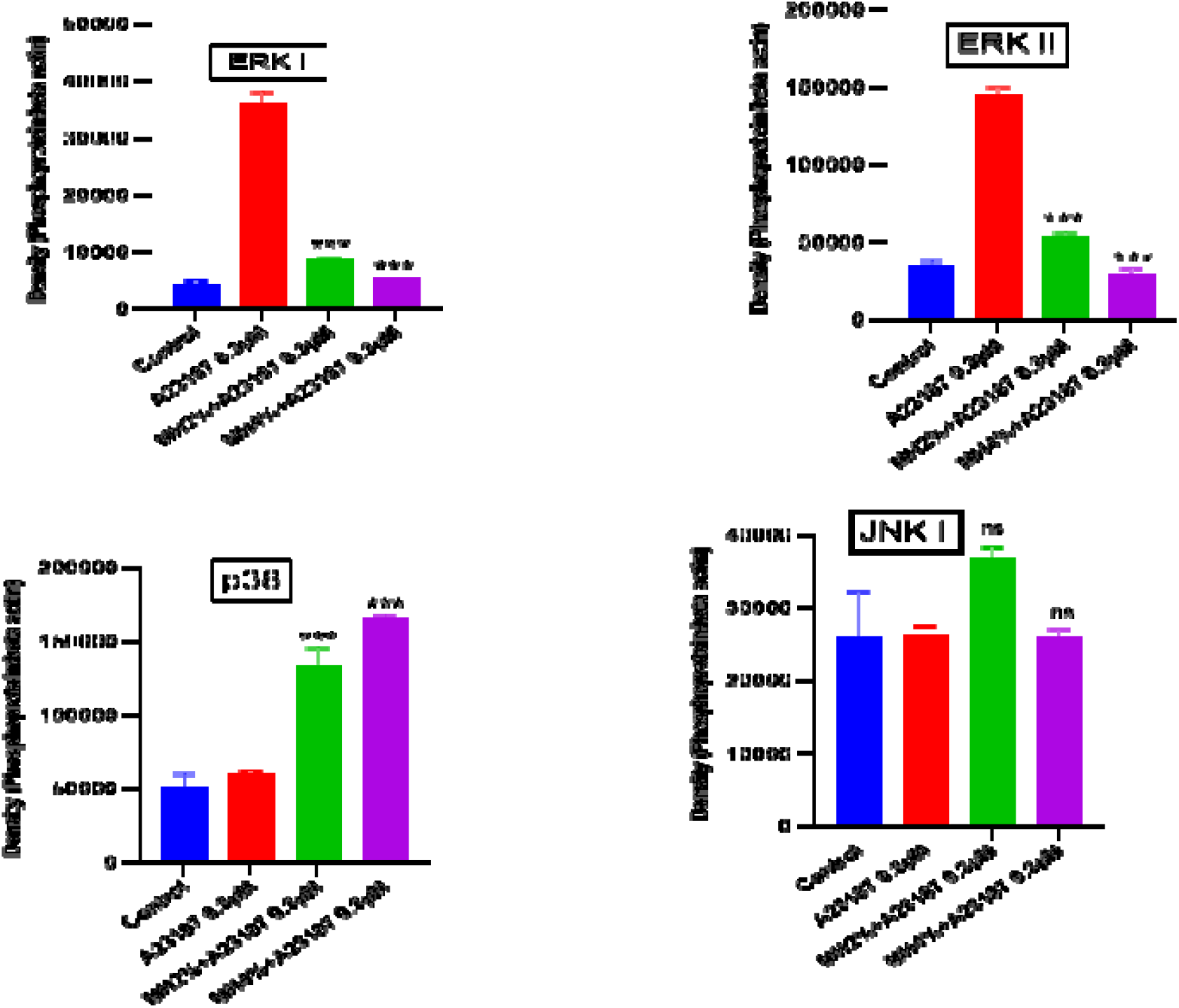
Quantification by densitometry of immunoblot showing the effect of MH on signalling MAPK. Densitometric analysis of immunoblot. Data is represented as analyses of protein densities of three different blots normalised to β-actin. Results are presented as Mean ± SEM. ***=0.000, non-significant in comparison to A23187 group.

## DISCUSSION

Mast cell degranulation and mediators afford a novel therapeutic target for mast cell related mastocytosis. We have in this study elucidated the effect of MGMH on mediator’s release and cell integrity. Our data suggested that 2% and 4% MGMH were well tolerated by LAD2 cells. Manuka honey is rich in complex sugars which confer osmotic properties at higher concentration and this may explain the reason for the cytotoxic actions seen with MGMH at 6% and 8% (Alvarez-Suarez et al. 2014).

We have in this work demonstrated that MGMH inhibits mast cell degranulation following A23187 stimulation. A23187 is a non-immunologic stimulus of mast cell exocytosis which is mediated by increasing intravessicular Ca^2+^ levels (Wightman et al. 2002). Histamine is a known mediator of inflammation and driver of pain in several diseases especially asthma and interstitial cystitis amongst others (Lamale et al. 2006). However, MGMH failed to inhibit its release despite histamine being a pre-formed mediator. This is an intriguing finding; suggesting that MGMH has differential actions in inhibiting mast cell degranulation. Both histamine and β-hexosaminidase are granular contents of mast cells, but histamine is stored with serglycin proteoglycan which has been demonstrated to show morphological and functional heterogeneity whilst β-hexosaminidase is of lysosomal origin (Wernersson and Pejler 2014). It might be that different types of granules are differentially sensitive to stimulated degranulation and this could account for selective effect of MGMH on mast cell degranulation observed in our study.

Pro-inflammatory cytokines particularly TNF-α, MCP-1 and IL-17A are increased in many inflammatory conditions compared to controls (Logadottir et al. 2014; Liu and Kuo 2015). However, these cytokines were not detected in our model. A plausible explanation for this could be related to the use of A23187; this is because cytoplasmic and nuclear granules respond differently to mast cell activating stimuli (Gaudenzio et al. 2016). With respect to other detectable pro-inflammatory cytokines; a strong and positive correlation was observed between the symptoms of IC and GM-CSF levels in a cyclophosphamide induced model of IC. This is in addition to its role as a promoter of the inflammatory response through the attraction of monocyte and granulocytes to sites of inflammation (Smaldone et al. 2009; Bhattacharya et al. 2015). Likewise, IL-8 is a chemotactic cytokine that attracts monocytes, lymphocytes, basophils and eosinophils to sites of inflammation (Turner et al. 2014). In addition, it is a surrogate marker in allergic asthma and Chronic Obstructive Pulmonary Disease (COPD) and bronchiectasis (Garth et al. 2018). Thus, inhibition of IL-8 by MGMH suggests possible potential in the treatment of these airway diseases.

The Mitogen Activated Protein Kinases (MAPK) are a diverse class of kinases that play a crucial role in signal transduction, neural plasticity and the inflammatory response. The members of this group include: extracellular signal-regulated kinase (ERK), p38 and c-Jun N-terminal kinase (JNK) (Ji et al. 2009). Activation of MAPK leads to gene transcription that regulates the synthesis of cytokines. For instance, ERK regulates the production of GM-CSF in human cultured mast cells and that could explain why the down-regulation of ERK by MGMH leads to inhibition of GM-CSF following A23187 challenge (Kimata et al. 2000). Similarly, MGMH increased the expression of p38 and as IL-1 and TNF-α are direct downstream targets of p38 (Kyriakis and Avruch 1996) this might explain why MGMH the slight increase in the production of IL-1β seen in our study.

Following Ca^2+^ influx in A23187 degranulation, membrane fusion leads to the formation of a SNARE complex which activates the PKC, RAS and subsequently the MAPK pathways (Suzuki and Verma 2008). Activation of the MAPK cascade has been well demonstrated in various models of hyperalgesia particularly in Bee venom, the formalin test and thermal models of pain (Cui et al. 2008; Ji et al. 2009). Furthermore, activation of ERK was well demonstrated in a cyclophosphamide induced model of IC in rats which correlated strongly with visceral hyperalgesia. Thus, ERK kinases are important targets of treatment. The fact that MGMH significantly suppressed the expression of ERK I and ERK II shows the potential benefit of this agent in IC and other inflammatory diseases.

## CONCLUSION

MGMH demonstrated a suppressive action on both the degranulation of mast cells and the release of IL-8 and GM-CSF. The latter action being mediated through the down regulation of ERK I and ERK II. This suggests that MGMH Honey could, potentially, be a lead compound in treatment of IC/PBS and other inflammatory disorders. Previous work from our group had demonstrated that MGMH offered significant inhibition of mast cell degranulation in comparison with currently available mast cell modulators used in the treatment of IC/PBS (Birch et al. 2011.

## ACKNOWLEDGEMENT

The study was partly supported by the Petroleum Technology Development Fund (PTDF), Nigeria.

